# Resting-state fMRI Signals of Intelligent People Wander in a Larger Space

**DOI:** 10.1101/529362

**Authors:** Aslan S Dizaji, Mohammad-Reza Khodaei, Hamid Soltanian-Zadeh

## Abstract

Natural intelligence is one of the vastly explored research areas in cognitive science. Its evolution and manifestation through behavioral patterns in animal kingdom have been extensively investigated. Since early days of cognitive sciences, there have been considerable efforts to simulate intelligent behaviors through high-level cognitive models. In the framework of the computational theory of mind, production systems are top-down models which simulate intelligent behaviors by invoking their behavioral manifestations. These models describe an intelligent behavior as structured mental programming which decomposes a complex task into simpler independent parts, each one represented by a cognitive enclosure where attention is sequentially devoted, and finally the information obtained from all cognitive enclosures is integrated to accomplish the task. In this article, we investigate the relations between these models of intelligence and resting-state fMRI signals. Based on these models, we hypothesize that the capacity of distinct mental representations is the core feature of intelligent behaviors. Therefore, we reason that resting-state fMRI signals of intelligent individuals wander in a larger space and can be divided to more well-separated independent components. This may be interpreted as the functional equivalence of one of the most celebrated structural correlates of intelligence, its positive association with the total brain volume. In the general framework of topological data analysis, using a well-established non-linear dimensionality reduction method, we show that indeed resting-state fMRI signals of intelligent individuals occupy a larger space and can be divided to more well-separated components with less connections in the reduced two-dimensional space. To our knowledge, this is the first attempt to relate the functional space of resting-state fMRI signals with the behavioral signatures of the human intelligence.

## INTRODUCTION

In an engaging story, Kevin Laland devotes one chapter of his book (Laland, 2018) to explain the evolution of intelligence and investigates systematically its relation with the brain size among animal kingdom. He starts by mentioning the “cultural drive hypothesis” which asserts that the proliferation of behavioral innovations by cultural transmission caused animals to exploit the environment in new ways, and thus increased the rate of genetic evolution. This suggests that the capability of animals to conceive novel solutions to surrounding challenges and to imitate the good ideas of other animals would offer individuals an advantage in the struggle to survive and reproduce. These abilities must have some substrate in the brain such as its volume. Assuming this, selection for ingenuity and social learning capabilities would favor brains with larger of this correlate, which in turn would further enhance their innovation and social learning. It is speculated that this cultural drive had culminated in humans as the most innovative and culturally dependent species. Extensive analysis of this hypothesis in primates, introduced the notion of “primate g” similar to the “g factor” or “general intelligence” in humans. “General intelligence” is a variable that summarizes positive correlations among different cognitive tasks, reflecting the fact that a performance on one type of cognitive tasks correlates to the performance on other kinds of cognitive tasks for the same person. Furthermore, it has been shown that both “primate g” and “general intelligence” have positive correlations with the total brain volume. To complement this line of argument, a question might be asked: is it possible to find some other brain correlates of intelligence which fit better with the depicted picture of the evolution of intelligence? Higher intelligence comes along with higher rates of innovation and social learning which both require a brain with more distinct mental representations devoted to each case of innovative or copied behavior.

Mental representation is the cornerstone of the computational theory of mind. The computational theory of mind asserts that human mind is an information processing system realized by the computation of neural activities to generate cognition. Computation is commonly understood in terms of Turing Machines which manipulate symbols according to a set of rules. The critical aspect of such a computational model is that it is independent from particular physical details of the machine that implements the computation and it requires mental representations in the form of symbols representing other objects. At the heart of the computational theory of mind is the idea that thoughts are a form of computation and a computation is by definition a systematic set of laws for the relations among representations (Pinker, 2009). Considering this theory, Steven Pinker defines intelligence as *“the ability to attain goals in the face of obstacles by means of decisions based on rational (truth-obeying) rules … intelligence consists of specifying a goal, assessing the current situation to see how it differs from the goal, and applying a set of operations that reduce the difference”* (Pinker, 2009). This is a first-principle definition intending to model the phenomenology of functional manifestation of intelligence at the course-grained scale (Bassett, Zurn, & Gold, 2018). Based on this definition, cognitive models of intelligence decompose cognition into meaningful functional elements which, independently from the details of biological implementations in the brain, perform cognitive tasks to explain behaviors (Kriegeskorte & Douglas, 2018). Production systems are such models providing a computational basis for intelligent behaviors. These are symbolic models using rules and logic and they operate on symbols rather than sensory signals. A “production” is a cognitive action triggered according to an if-then rule. A set of such rules specifies the conditions under which each of a range of productions is to be executed. The conditions refer to current goals and knowledge in memory. The actions can modify the internal state of goals and knowledge. A model specified using this formalism will generate a sequence of productions, which might resemble our conscious stream of thought while working toward some cognitive goal (Kriegeskorte & Douglas, 2018). Here, the main challenge is to relate these cognitive models of intelligence formalized in task-performing computational models to the experimental data of “general intelligence” provided by brain images and psychometric tests.

The notion of “general intelligence” in humans, discussed above, was first introduced in the early 20th century by Charles Spearman who discovered that performance measures in diverse cognitive tests are positively correlated with each other; in other words, to some extent, the same individual performs similarly in very different tasks. To explain this result, Spearman proposed the hypothesis of a “g factor” or “general intelligence” which makes contribution to the success in diverse forms of cognitive tasks (Deary, Penke, & Johnson, 2010; Duncan et al., 2000; Hampshire, Highfield, Parkin, & Owen, 2012). Through neuroimaging experiments, it has been shown that a focal region of brain called multiple-demand cortex, which lies at the parietal and frontal cortices, is activated for a range of different cognitive tests requiring high “general intelligence” (Duncan, 2010b; Fedorenko, Duncan, & Kanwisher, 2013). “General intelligence” is further divided to two subcategories: crystalized intelligence and fluid intelligence, where crystalized intelligence refers to the knowledge obtained through education and experience, and fluid intelligence points to the more abstract reasoning and problem-solving abilities including adaption of acquired knowledge to a complex environment (Duncan, 2010a). In psychometrics, fluid intelligence can be measured by a test like Raven's Progressive Matrices and John Duncan describes it by invoking the cognitive models of intelligence through structured mental programming of production systems (Duncan, 2010a; Duncan, 2010b): in any goal-directed behavior, the task at hand is divided robustly into independent sub-problems and attention focuses on these sub-problems sequentially to provide solution to the original problem. Furthermore, for each sub-problem, a separate cognitive enclosure is devoted, which consists of similar elements around some task-relevant features, and the transitions from one cognitive enclosure to another one are managed by corresponding transitions among distributed and independent patterns of activity in the multiple-demand cortex. As a result, the procedure of structured mental programming is to reduce the compositionality or complexity of a task through decomposing it into simpler separately attended parts representing each one of them by a cognitive enclosure and then integrating the information obtained from all of them to accomplish the task (Duncan, Chylinski, Mitchell, & Bhandari, 2017; Tschentscher, Mitchell, & Duncan, 2017). We believe, any attempt to relate the psychometric measures of fluid intelligence to the features of brain images like fMRI should consider this high-level description.

However, recent trend in the literature using both neuroimaging and psychometric tests is rarely considering these high-level cognitive models of intelligence. The neural efficiency hypothesis (Neubauer & Fink, 2009) is one of the early ad hoc models of intelligence which states that more intelligent individuals use less brain resources when confronting with cognitively challenging tasks. The parieto-frontal integration theory (P-FIT) of intelligence (Basten, Hilger, & Fiebach, 2015; Jung & Haier, 2007) is another mainstream model which proposes that large-scale brain networks connecting brain regions within parietal and frontal cortices, underlie the biological basis of human intelligence. As it is evident, there is no reference to the definition of intelligent behaviors by structured mental programming of production systems in these two models. More recently, an emerging trend in the literature which aims to relate the characteristics of brain networks to intelligence is culminated to the network neuroscience theory of human intelligence (Barbey, 2018). This theory asserts that intelligence depends on the dynamic reorganization of brain networks which modifies its topology in the service of system-wide flexibility and adaptation. In this context, crystallized intelligence utilizes easy-to-reach network states that access prior knowledge on systems and experience, while fluid intelligence demands difficult-to-reach network states that support cognitive flexibility and adaptive problem-solving (Barbey, 2018). For example, (Schultz & Cole, 2016) show that individuals with higher intelligence reconfigure their resting-state fMRI networks less but more efficiently than the individuals with lower intelligence during cognitively demanding tasks. As another example, (Genç et al., 2018) show that branching of dendrites in the brain of intelligent individuals shows more efficient and sparse organization compared to less intelligent individuals. Also, there are other network models which use graph theoretical metrics to analyze functional connectivity of resting-state fMRI (Cole, Yarkoni, Repovs, Anticevic, & Braver, 2012; Song et al., 2008; M. P. van den Heuvel, Stam, Kahn, & Hulshoff Pol, 2009) to show that brain functional networks, specifically in the parietal and frontal cortices, have small world architecture, possessing high clustering coefficient (local efficiency) and short characteristic path length (global efficiency) (for null results, see (Kruschwitz, Waller, Daedelow, Walter, & Veer, 2018)). Furthermore, there are other data-driven models in the framework of network neuroscience (Bassett et al., 2018) which try to predict intelligence through its distributed representation by functional connectivity among many nodes of resting-state fMRI (Dubois, Galdi, Paul, & Adolphs, 2018; Finn et al., 2015; Greene, Gao, Scheinost, & Constable, 2018). What is missing in all these approaches is the direct link between the definition of intelligence or its characteristics captured by psychometric tests with the functional neuroimaging data. Any model of intelligence, either in the framework of network neuroscience (Bassett et al., 2018) or not, should determine its position in relation to the behavioral manifestation of intelligence implied in its definition by the process of structured mental programming of production systems. As wisely said, *“a brain map, at whatever scale, does not reveal the computational mechanism … the challenge ahead is to build computational models of brain information processing that are consistent with brain structure and function and perform complex cognitive tasks”* (Kriegeskorte & Douglas, 2018). We believe network neuroscience without referring to the theory-driven high-level cognitive models of intelligence, lacks the deep insight and might hinder the progress in the direction of deciphering the computational nature of intelligence.

In this article, by invoking the high-level cognitive model of intelligence implied in its definition, we hypothesized that one plausible correlate of intelligence could be the size of functional space in which the brain fMRI signals of an individual occupy. This is roughly commensurate with the capability of mental representations of distinct states in the brain. Moreover, the functional networks of resting-state fMRI are relatively similar to the functional networks of task-based fMRI suggesting that resting-state fMRI oscillations may reflect ongoing functional communication among brain regions (Deco, Jirsa, & McIntosh, 2011; Martijn P. van den Heuvel & Hulshoff Pol, 2010). For this reason, we used the high-quality resting-state fMRI data of Human Connectome Project (HCP). We first applied a mask to divide the voxels of multiple-demand cortex of brain to homogenous nodes. Then, in the general framework of topological data analysis (Lum et al., 2013; Saggar et al., 2018), we used a well-established non-linear dimensionality reduction technique, t-distributed Stochastic Neighbor Embedding (t-SNE) (Maaten & Hinton, 2008), to reduce the high-dimensional resting-state fMRI data to two-dimensions. Afterwards, by constructing the simplicial graph of the data for each subject, we found that the resting-state fMRI signals of intelligent people in the reduced two-dimensional space comprised of more nodes and thus occupy a larger space. Moreover, to further test the model of sequential mental programming of production systems, we hypothesized that distinct mental representations in the brain of intelligent individuals might be more separated from each other. We found some evidence for this hypothesis which indicates that the resting-state fMRI signals of intelligent people in the reduced two-dimensional space can be divided into more well-separated cognitive enclosures with less connections. Therefore, each one of these cognitive components potentially could be assigned to an independent sub-task of a decomposed complex task (Duncan et al., 2017; Tschentscher et al., 2017).

## MATERIALS AND METHODS

### HCP Data

In this article, the minimally preprocessed resting-state fMRI data of “100 Unrelated Subjects” were used. Both sessions of data (Resting State fMRI 1 Preprocessed & Resting State fMRI 2 Preprocessed) were downloaded from “ConnectomeDB” website (https://db.humanconnectome.org). We used this data which excludes family relations because we wanted to have a sample representative of the general population. The sample consists of 54 females and 46 males with the age range of 22–36+.

Acquisition parameters and minimal preprocessing of the resting-state fMRI data were described elsewhere (Glasser et al., 2013). Briefly, each subject underwent two sessions of resting-state fMRI on separate days, each session with two separate 14 min 33 s acquisitions generating 1200 volumes (customized Siemens Skyra 3 Tesla MRI scanner, TR = 720 ms, TE = 33 ms, flip angle = 52^º^, voxel size = 2 mm isotropic, 72 slices, matrix = 104 * 90, FOV = 208 * 180 mm, multiband acceleration factor = 8). The two runs acquired on the same day differed in the phase-encoding direction, left-right and right-left. In the main text of this article, we used the left-right run of the first session as a training-set and the left-right run of the second session as a validation-set for the parameters of t-SNE method and topological data analysis. In the supplementary material of this article, the same results were brought for the right-left run of the first and second sessions. Moreover, the HCP minimal preprocessing pipeline, which is used in this article, includes artifact removal, motion correction, and registration to the standard space.

To focus on the relevant regions of brain which coordinate intelligent behaviors, the mask of multiple-demand cortex (Fedorenko et al., 2013) was applied on the resting-state fMRI. Then to reduce dimensionality and also spatially smooth the data, the Shen 268-node atlas (Shen, Tokoglu, Papademetris, & Constable, 2013) was applied on the voxels of multiple-demand cortex of the resting-state fMRI to parcellate it to 132 functionally homogenous nodes (here, a node is the average signal of multiple neighboring voxels in the brain space). Additionally, we can interpret each time-point of the resting-state fMRI as a point in the 132-dimensional space which was subsequently mapped to two-dimensions by t-SNE method. After performing topological data analysis, the data was converted to a simplicial graph in two-dimensions which is comprised of many nodes and edges (here, a node is the average position of multiple neighboring time-points in the reduced two-dimensional space).

In the HCP data-set, fluid intelligence was quantified using matrix reasoning tests. The 24-item (3 bonus) Penn Matrix Reasoning Test (PMAT), a test of non-verbal reasoning ability that can be administered in under 10 min, was used. The PMAT is designed to parallel many of the psychometric properties of the Raven’s Standard Progressive Matrices test, while limiting learning effects and expanding the representation of the abstract reasoning construct (Dubois et al., 2018; Greene et al., 2018). The scores range for “100 Unrelated Subjects” is 6-24.

### t-SNE Method

One main step of topological data analysis is to reduce the dimensionality of data. For this purpose, we used t-SNE method which is a non-linear dimensionality reduction technique allows for preservation of the “local” structure in the original high-dimensional space after projection into the low-dimensional space (Maaten & Hinton, 2008). This is usually not possible with classical linear methods such as principal component analysis (PCA) or multidimensional scaling (MDS). We bring here a brief description of t-SNE method and refer interested readers to the original publication.

t-SNE is a non-linear dimensionality reduction method which converts the high-dimensional data *X* = {*x*_*1*_, *x*_*2*_, *x*_*3*_,…, *x*_*n*_} into two-dimensional map *Y* = {*y*_1_, *y*_*2*_, *y*_3_,…, *y*_*n*_} that can be shown in athat can be shown in a scatterplot. The aim of dimensionality reduction is to preserve as much of the significant structure of the high-dimensional data as possible in the low-dimensional map. t-SNE is capable of both capturing the local and global structures of the high-dimensional data.

In the high-dimensional space, the distances (which are measured based on a metric specified by the user) are converted into probabilities using a Gaussian distribution. The similarity of data-point *x*_*j*_ data-point *x*_*i*_ is the conditional to probability *p_j\i_*, that *x_*i*_* would pick *x*_*j*_ as its neighbor, if neighbors were picked in proportion to their probability density under a Gaussian centered at *x*_*i*_. The joint probability defined as symmetrized conditional probabilities is used for this purpose:

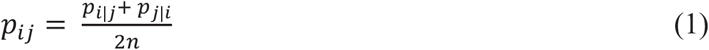

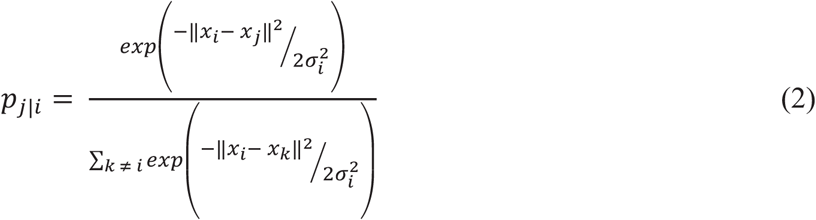

The variance *σ*_*i*_ of the Gaussian that is centered over each high-dimensional data-point, *x*_*i*_, should be determined beforehand. For this purpose, t-SNE performs a binary search for the value of *σ*_*i*_ that produces a *P*_*i*_ with a fixed perplexity specified by the user. The perplexity is defined as:

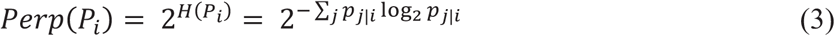

The perplexity can be interpreted as a smooth measure of the effective number of neighbors. The performance of t-SNE is fairly robust to changes in the perplexity, whose typical values are between 5 and 50.

In the low-dimensional map, a Student t-distribution with one degree of freedom (which is the same as a Cauchy distribution) is used that has much heavier tails than a Gaussian to convert distances (which are again measured based on a metric specified by the user) into probabilities. The joint probability is used for this purpose:

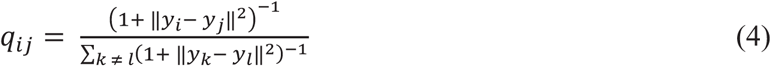

If the map points *y*_*i*_ and *y*_*j*_ correctly model the similarity between the high-dimensional data-points *x*_*i*_ and *x*_*j*_, the joint probabilities *p*_*ij*_ and *q*_*ij*_ will be equal. Based on this observation, t-SNE aims to find a low-dimensional data representation that minimizes the discrepancy between the distributions *p*_*ij*_ and *q*_*ij*_. A natural measure of the dissimilarity between *p*_*ij*_ and *q*_*ij*_ is the Kullback-Leibler divergence which should be minimized between a joint probability distribution, *P*, in the high-dimensional space and a joint probability distribution, *Q*, in the low-dimensional space using a gradient descent method. The cost function *C* for this minimization (which is non-convex and thus in each run using the same optimization parameters and starting from a random initial condition does not necessarily converge to the same minimum) is given by:

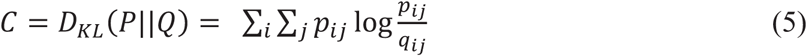

### Topological Data Analysis

The topological data analysis seeks to understand the “shape” of data instead of focusing on pairwise relationships as the fundamental building block of more traditional data mining approaches. In the topological data analysis, topology acts as a geometric approach to recognize patterns within data, thus it is able to discover insights about the meaningful structures of data. Moreover, there are three key concepts of topological methods, coordinate freeness, invariance to deformation, and compressed representations of shapes which are of particular value for applications to data analysis. The topological data analysis has generally three inputs: the filter function (a real-valued function which maps high-dimensional data-points to two-dimensional data-points and represents similar points nearby than dissimilar points; also, it may independently require a few inputs), and two binning parameters (“resolution” (R) which determines the number of bins in each dimension of the two-dimensional space, and “percent overlap” or “gain” (G) which determines the percentage of overlapped area between two consecutive bins). Afterwards, the method constructs a network of nodes with edges between them named simplicial graph which captures the overall shape of data. The coordinates of any individual node have no particular meaning and only the relations of nodes to each other bear meaning. The nodes represent sets of data-points and two nodes are connected if and only if their corresponding aggregation of data-points have a point in common (Lum et al., 2013).

The topological data analysis was applied successfully on task-based fMRI to reveal the dynamical organization of the brain with high temporal resolution (Saggar et al., 2018). In this article, we adopted a modified version of this method briefly described in the following paragraph in five steps. Also, we performed a data-driven optimization for extensive range of four input parameters (t-SNE parameters: distance metric and perplexity; binning parameters: resolution and gain) to show that the presented results are stable.

At the first step, each subject’s parcellated fMRI data were transformed into a 2D matrix, such that the rows correspond to the individual time-points (or volumes) and the columns correspond to the intensity value at each homogenous node of the brain. Thus, each row of this matrix represents the entire brain volume at any time-point during the session. For the HCP data-set, the size of each individual’s 2D input matrix was [1200 × 132]. At the second step, t-SNE (as it is implemented in the built-in function of MATLAB 2017b) was used as a filtering function for dimensionality reduction to project the high-dimensional data (in 132 dimensions) to two dimensions. This filtering was done several times to find the best two-dimensional map with the minimum value of cost function *C*, also to avoid the presence of outliers adversely effects the shape of the data. This means that we do not let each one of the filtered time-points in the two-dimensional space being distant from the rest of the time-points more than one bin size. This generates fair comparison of functional space across subjects. The default values of all input parameters of t-SNE function were used except the two main parameters, distance metric and perplexity. These two parameters were perturbed extensively to show the validity of results as it will be explained in the Results Section. At the third step, to encapsulate the two-dimensional representation generated by the filtering step, binning was employed. The binning step partitions the two-dimensional space between the minimum and maximum values of time-points in each dimension into overlapping bins by using two parameters, number of bins (or resolution (R)) and percentage of overlap between consecutive bins (or gain (G)), and then combines all time-points inside a bin to a single node. This kind of binning again provides fair comparison of functional space across subjects, but at the same time generates relatively conservative condition to test our hypotheses with the graph theoretical metrics. Again, at this step, a wide-range of values for two parameters, R and G, were used to test the reliability of our results for both training and validation data-sets as it will be explained in the Results Section. At the fourth step, to generate a simplicial graph from the two-dimensional compressed representation, the nodes which are the average of all time-points inside overlapping bins were connected with edges if they shared time-points. Finally, at the fifth step, three relevant graph theoretical metrics, the number of nodes, the number of connected components, and the average degree centrality of the simplicial graph were calculated for each subject and subsequently correlated with intelligence across all subjects.

## RESULTS

The main hypothesis of this article is that the functional space of resting-state fMRI signals of more intelligent people is larger compared to less intelligent individuals. This hypothesis is originated from the observation that intelligent individuals have higher capacity for mental representations of different states (in this paper, we measure intelligence or intelligence quotient (IQ) by PMAT score which quantifies fluid intelligence). To test this hypothesis, the topological data analysis was performed on resting-state fMRI to map the high-dimensional data to two-dimensions, so to be able to extract the shape of data. Fig. 1 depicts schematically the five steps of this method which we used in this paper: constructing the input matrix (Fig. 1 (a)), filtering by t-SNE (Fig. 1 (b)), binning (Fig. 1 (c)), constructing the simplicial graph (Fig. 1 (d)), and calculating the three graph theoretical metrics (Fig. 1 (e)). It was expected that the shape of two-dimensional map of resting-state fMRI does not have any specific structure. However, to test our main hypothesis, it is possible to define minimally the following graph theoretical metric: the number of nodes of the two-dimensional simplicial graph of each subject which roughly represents the area of functional space of resting-state fMRI signals in the two-dimensional space and be expected to correlate positively with intelligence. This is a fair comparison across subjects but relatively conservative measure due to our implementation of topological data analysis. Moreover, the higher-level cognitive model of intelligence explains an intelligent behavior by sequential mental programming of production systems and states that any intelligent behavior consists of decomposing a complex task to simpler independent parts, encompassing each one of them by a well-separated cognitive enclosure, and finally, integrating the information obtained from all of them to accomplish the task. To test this model, two further graph theoretical metrics were calculated. First, the number of connected components of the simplicial graph of each subject which roughly represents the number of independent well-separated cognitive enclosures was calculated and expected to positively associate with intelligence. Second, the average degree centrality of the simplicial graph of each subject which roughly represents the connectedness of cognitive enclosures to each other was calculated and expected to correlate negatively with intelligence. Fig. 2 shows the application of the main three steps of topological data analysis (steps two, three, and four) on two cases of representative subjects with low IQ (subject number = 100408, IQ = 7) on panels (a), (b) and (c) and high IQ (subject number = 129028, IQ = 24) on panels (d), (e), and (f). The data is from the left-right run of the second session of HCP data-set and the same values were used for the four input parameters of topological data analysis (distance metric = chebychev, perplexity = 50, resolution = 15, gain = 6). The obtained number of nodes, number of connected components, and average degree centrality for low IQ person are 128, 25, and 1.94 respectively. For the high IQ person, the corresponding values are 167, 51, and 1.62 respectively. As it is clear, the number of nodes and number of connected components are higher for high IQ person compared to low IQ person while the average degree centrality is higher for low IQ person compared to high IQ person, all in accordance with our three hypothesis.

**Fig. 1:**
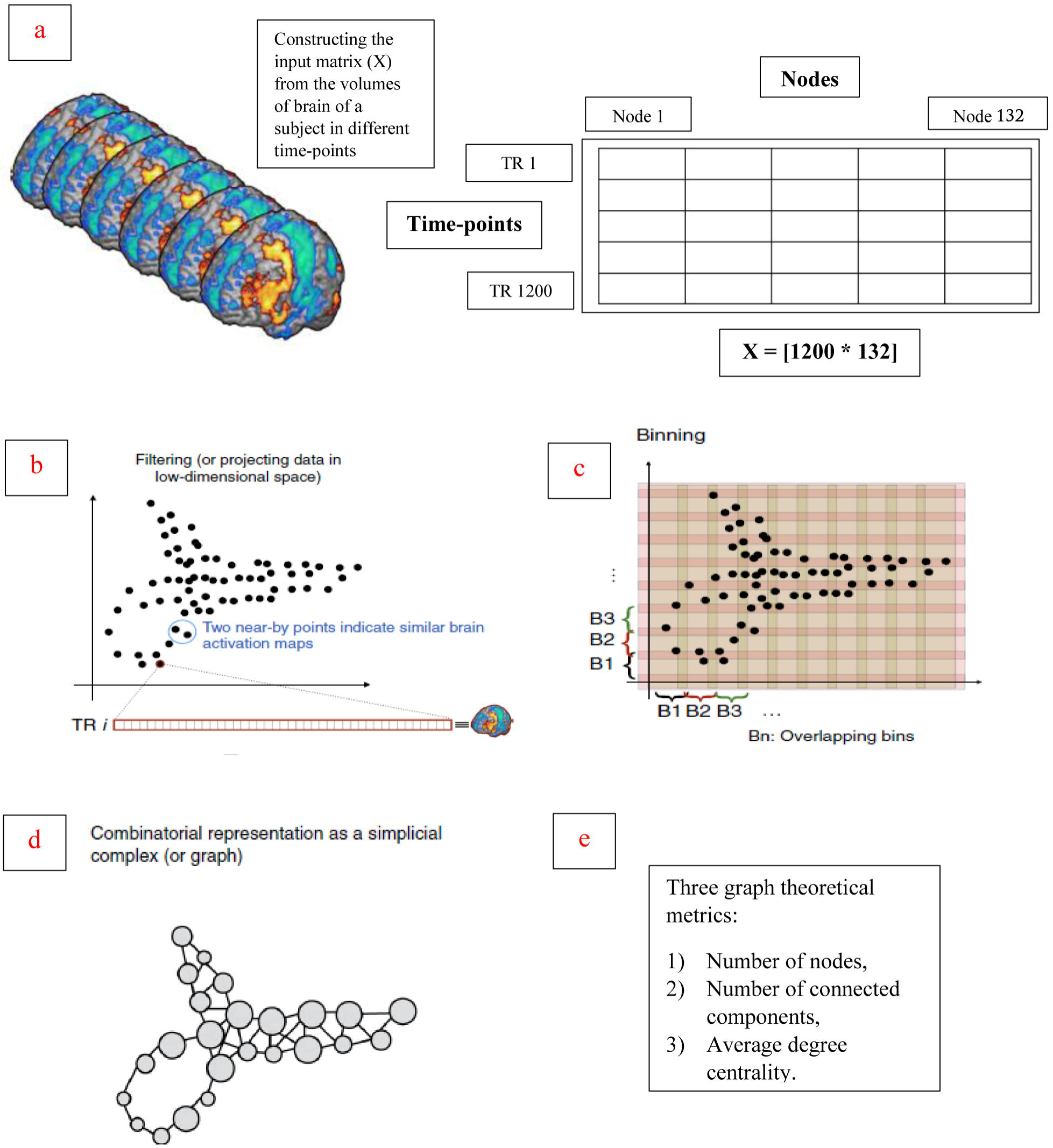
A schematic figure showing the five steps of the topological data analysis method as it was implemented in this paper and described with more details in the Materials and Methods Section: **(a)** Constructing the input matrix, **(b)** Filtering by t-SNE, **(c)** Binning, **(d)** Constructing the simplicial graph, **(e)** Calculating the three graph theoretical metrics. Parts of the figure were reproduced from the Fig. 1 of (Saggar et al., 2018).

**Fig. 2:**
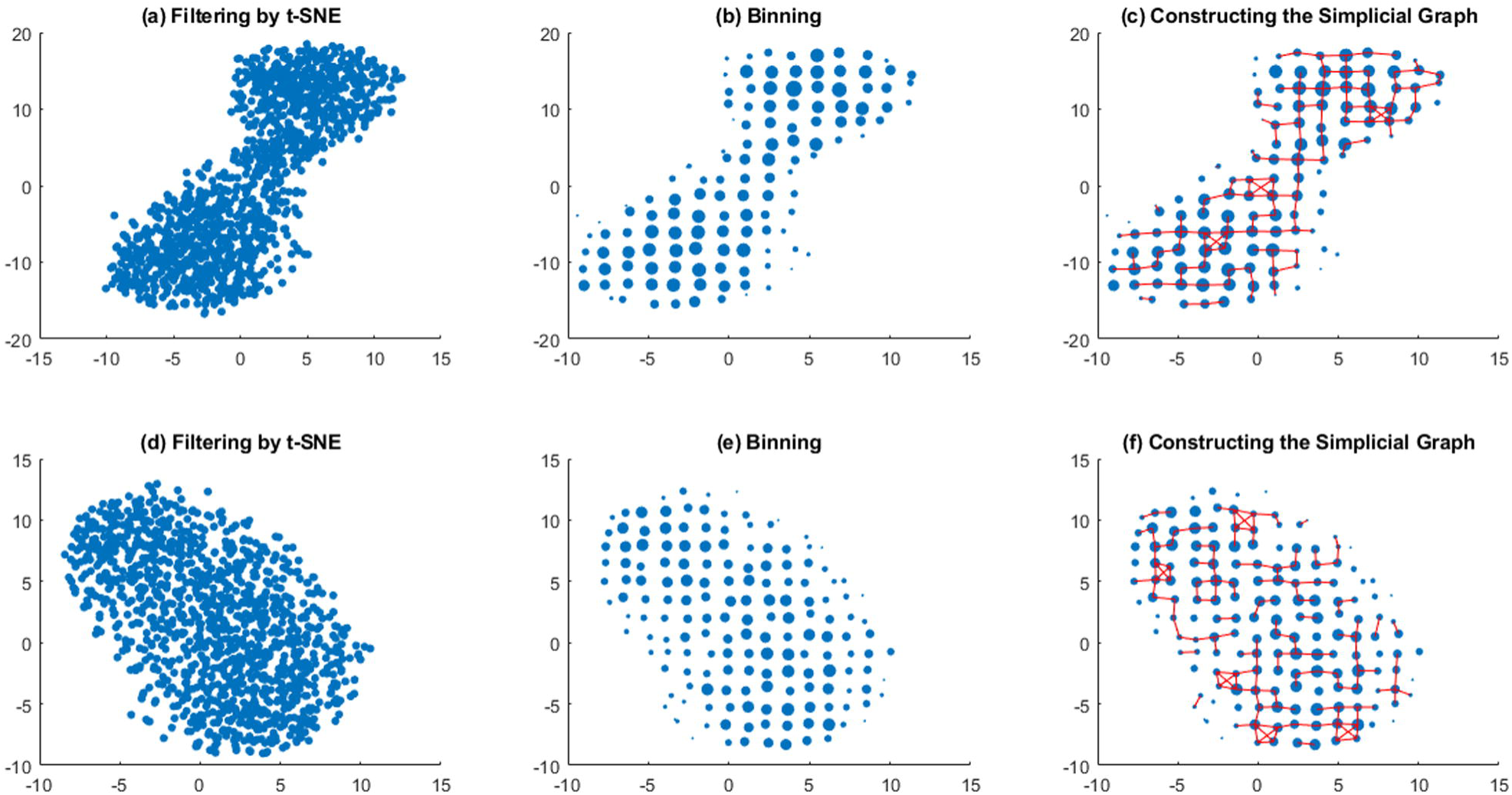
The application of the main three steps of topological data analysis (steps two, three, and four) on two cases of representative subjects with low IQ (subject number = 100408, IQ = 7) on panels **(a)**, **(b)** and **(c)** and high IQ (subject number = 129028, IQ = 24) on panels **(d)**, **(e)**, and **(f)**. The data is from the left-right run of the second session of HCP data-set and the same values were used for the four input parameters of topological data analysis (distance metric = chebychev, perplexity = 50, resolution = 15, gain = 6). The obtained number of nodes, number of connected components, and average degree centrality for the low IQ individual are 128, 25, and 1.94 respectively. For the high IQ individual, the corresponding values are 167, 51, and 1.62 respectively. As it is clear, the number of nodes and number of connected components are higher for high IQ person compared to low IQ person while the average degree centrality is higher for low IQ person compared to high IQ person, all in accordance with our three hypothesis.

Furthermore, to test the reliability of our results for different sets of input parameters of the topological data analysis (t-SNE: distance metric and perplexity; binning: resolution and gain), the method was run for a wide-range of parameters. Fig. S1 depicts the resultant simplicial graphs for different parameters of t-SNE, distance metric (euclidean, cityblock, chebychev, correlation, spearman) and perplexity (10, 20, 30, 40, 50), while using the fixed values for two binning parameters (resolution = 20 and gain = 6) for a representative subject. Additionally, Fig. S2 shows the resultant simplicial graphs for different parameters of binning, resolution (10, 15, 20, 25, 30) and gain (2, 4, 6, 8, 10), while using the fixed values for two filtering parameters (distnace metric = chebychev and perplexity = 30) for a representative subject. As it is clear from Fig. S1 and Fig. S2, the resultant simplicial graphs of resting-state fMRI change considerably across different parametes but do not have any particular structure as it is the case in the task-based fMRI (Saggar et al., 2018).

To systematically investigate the relationships of the three graph theoretical metrics of the simplicial graph of each subject with intelligence, we followed the following procedure:

1. The topological data analysis was performed on the resting-state fMRI of the left-right run of the first session of HCP data-set across all subjects for a range of two filtering parameters, distance metric (euclidean, cityblock, chebychev, correlation, spearman) and perplexity (10, 20, 30, 40, 50), while using the fixed values of two binnig parameters, (resolution = 20 and gain = 6). Then three graph theoretical metrics - the number of nodes, the number of connected componenets, and the average degree centrality of the simplicial graph of each subject – were calculted and correlated with intelligence across subjects. Fig. 3 summarizes these results by showing the correlation values (Fig. 3 (a) (d) (g)) and their corresponding p-values (Fig. 3 (b) (e) (h)). All the correlation values are positive for the first and second metrics while most of them are negative for the third metric in accordance with our hypotheses. Fig. 3 (c) shows the regression plot for the lowest p-value (0.03) and highest correlation (r = 0.22) between the number of nodes of simplicial graphs and intelligence across subjects. Moreover, Fig. 3 (f) shows the regression plot for the lowest p-value (0.012) and highest correlation (r = 0.25) between the number of connected components of simplicial graphs and intelligence across subjects. Finally, Fig. 3 (i) shows the regression plot for the lowest p-value (0.038) and highest absolute correlation (r = −0.21) between the average degree centrality of simplicial graphs and intelligence across subjects.
2. The sets of parameters which generate p-values less than 0.1 for the correlation of the first graph theoretical metric - the number of nodes - with intelligence, were selected to further investigate the effects of binning parameters, resolution and gain. There are five sets of those parameters in the pairs of (distance metric, perplexity): (chebychev, 20), (chebychev, 30), (chebychev, 50), (correlation, 40), and (spearman, 30). The topological data analysis was performed once again on the resting-state fMRI of the left-right run of the first session of HCP data-set across all subjects for each one of these sets of two filtering parameters, while perturbing the values of two binning parameters, resolution (10, 15, 20, 25, 30) and gain (2, 4, 6, 8, 10). All the correlation values are positive for the first metric, while most of them are positive for the second metric, and most of them are negative for the third metric in accordance with our hypotheses. The best set of two filtering parameters with lower p-values across all three metrics was (spearman, 30). Fig. 4 summarizes the results obtained from this set in a wide-range of values of resolution (10, 15, 20, 25, 30) and gain (2, 4, 6, 8, 10) parameters by showing the correlation values (Fig. 4 (a) (d) (g)) and their corresponding p-values (Fig. 4 (b) (e) (h)). Fig. 4 (c) shows the regression plot for the lowest p-value (0.04) and highest correlation (r = 0.21) between the number of nodes of simplicial graphs and intelligence across subjects. Moreover, Fig. 4 (f) shows the regression plot for the lowest p-value (0.042) and highest correlation (r = 0.28) between the number of connected components of simplicial graphs and intelligence across subjects. Finally, Fig. 4 (i) shows the regression plot for the lowest p-value (0.001) and highest absolute correlation (r = −0.32) between the average degree centrality of simplicial graphs and intelligence across subjects.
3. The same five sets of filtering parameters were selected ((chebychev, 20), (chebychev, 30), (chebychev, 50), (correlation, 40), and (spearman, 30)) to run the topological data analysis on the resting-state fMRI of the left-right run of the second session of HCP data-set across all subjects to further investigate the generalization of our results. For each one of these sets of two filtering parameters, two binning parameters were purtubed - resolution (10, 15, 20, 25, 30) and gain (2, 4, 6, 8, 10). All the correlation values are positive for the first metric, while most of them are positive for the second metric, and most of them are negative for the third metric in accordance with our hypotheses. The best set of two filtering parameters with lower p-values across all three metrics was (chebychev, 50). Fig. 5 summarizes the results obtained from this set in a wide-range of values of resolution (10, 15, 20, 25, 30) and gain (2, 4, 6, 8, 10) parameters by showing the correlation values (Fig. 5 (a) (d) (g)) and their corresponding p-values (Fig. 5 (b) (e) (h)). Fig. 5 (c) shows the regression plot for the lowest p-value (0.0052) and highest correlation (r = 0.28) between the number of nodes of simplicial graphs and intelligence across subjects. Moreover, Fig. 5 (f) shows the regression plot for the lowest p-value (0.00061) and highest correlation (r = 0.34) between the number of connected components of simplicial graphs and intelligence across subjects. Finally, Fig. 5 (i) shows the regression plot for the lowest p-value (0.00081) and highest absolute correlation (r = −0.33) between the average degree centrality of simplicial graphs and intelligence across subjects.

**Fig. 3:**
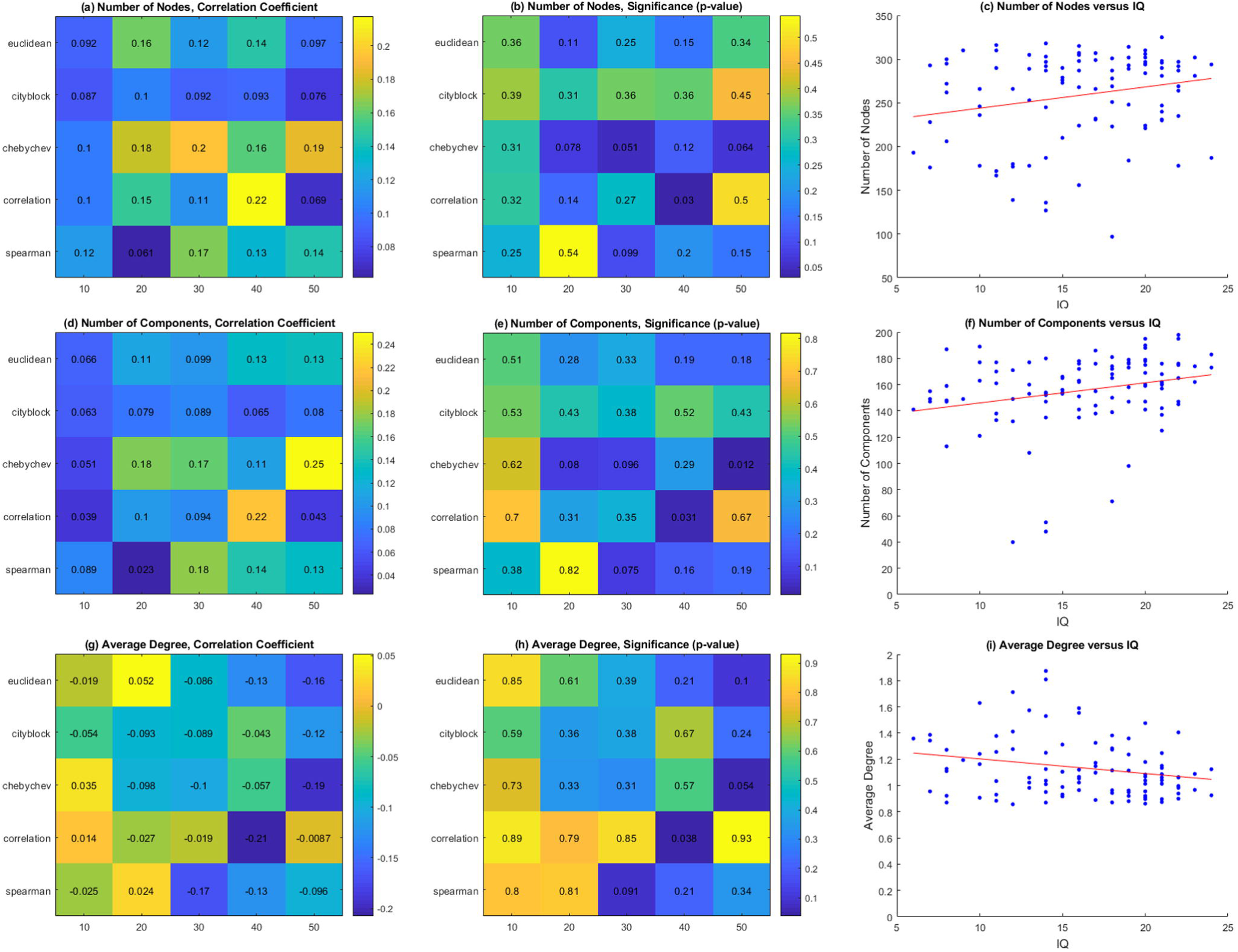
The number of nodes, number of connected components, and average degree centrality of simplicial graphs of all subjects (data is from the left-right run of the first session of HCP data-set) were calculated and then correlated with intelligence for a range of two input parameters of t-SNE function, distance metric (euclidean, cityblock, chebychev, correlation, spearman) and perplexity (10, 20, 30, 40, 50), using fixed values of two input parameters of binning, resolution = 20 and gain = 6. This was performed to show that the number of nodes and number of connected components associate positively with intelligence while the average degree centrality correlates negatively with intelligence across a wide-range of parameters. **(a) (b)** Correlation coefficients and their corresponding p-values for the number of nodes of simplicial graphs for a range of values of distance metric (y axis) and perplexity (x axis). **(c)** The number of nodes versus intelligence (IQ) for the lowest p-value (0.03) and highest correlation (r = 0.22). **(d) (e)** Correlation coefficients and their corresponding p-values for the number of connected components of simplicial graphs for a range of values of distance metric (y axis) and perplexity (x axis). **(f)** The number of connected components versus intelligence (IQ) for the lowest p-value (0.012) and highest correlation (r = 0.25). **(g) (h)** Correlation coefficients and their corresponding p-values for the average degree centrality of simplicial graphs for a range of values of distance metric (y axis) and perplexity (x axis). **(i)** The average degree centrality versus intelligence (IQ) for the lowest p-value (0.038) and highest absolute correlation (r = −0.21).

**Fig. 4:**
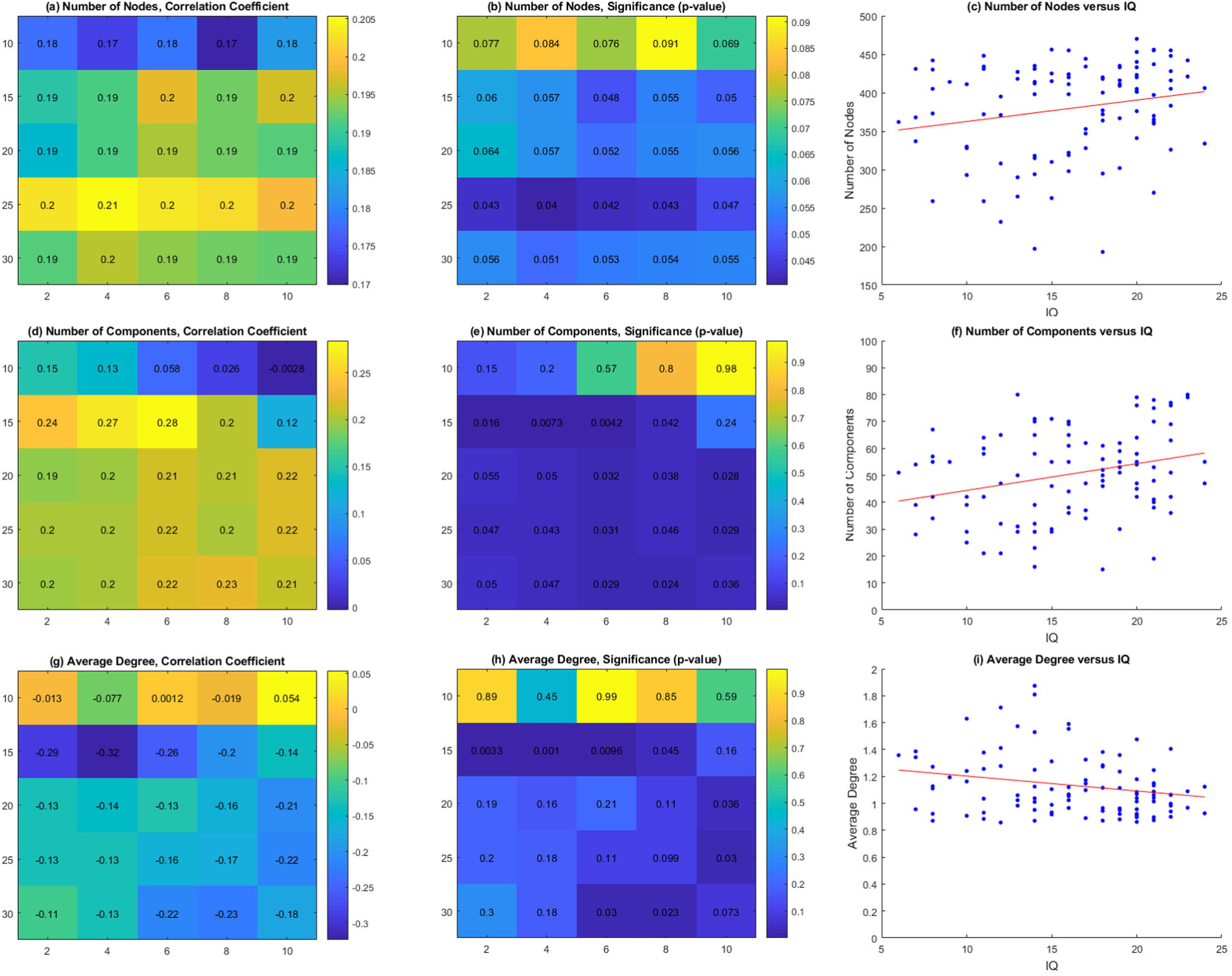
The number of nodes, number of connected components, and average degree centrality of simplicial graphs of all subjects (data is from the left-right run of the first session of HCP data-set) were calculated and then correlated with intelligence for a range of two input parameters of binning, resolution (10, 15, 20, 25, 30) and gain (2, 4, 6, 8, 10), using fixed values of two input parameters of t-SNE function, distance metric = spearman and perplexity = 30. This was performed to show that the number of nodes and number of connected components associate positively with intelligence while the average degree centrality correlates negatively with intelligence across a wide-range of parameters. **(a) (b)** Correlation coefficients and their corresponding p-values for the number of nodes of simplicial graphs for a range of values of resolution (y axis) and gain (x axis). **(c)** The number of nodes versus intelligence (IQ) for the lowest p-value (0.04) and highest correlation (r = 0.21). **(d) (e)** Correlation coefficients and their corresponding p-values for the number of connected components of simplicial graphs for a range of values of resolution (y axis) and gain (x axis). **(f)** The number of connected components versus intelligence (IQ) for the lowest p-value (0.042) and highest correlation (r = 0.28). **(g) (h)** Correlation coefficients and their corresponding p-values for the average degree centrality of simplicial graphs for a range of values of resolution (y axis) and gain (x axis). **(i)** The average degree centrality versus intelligence (IQ) for the lowest p-value (0.001) and highest absolute correlation (r = −0.32).

**Fig. 5:**
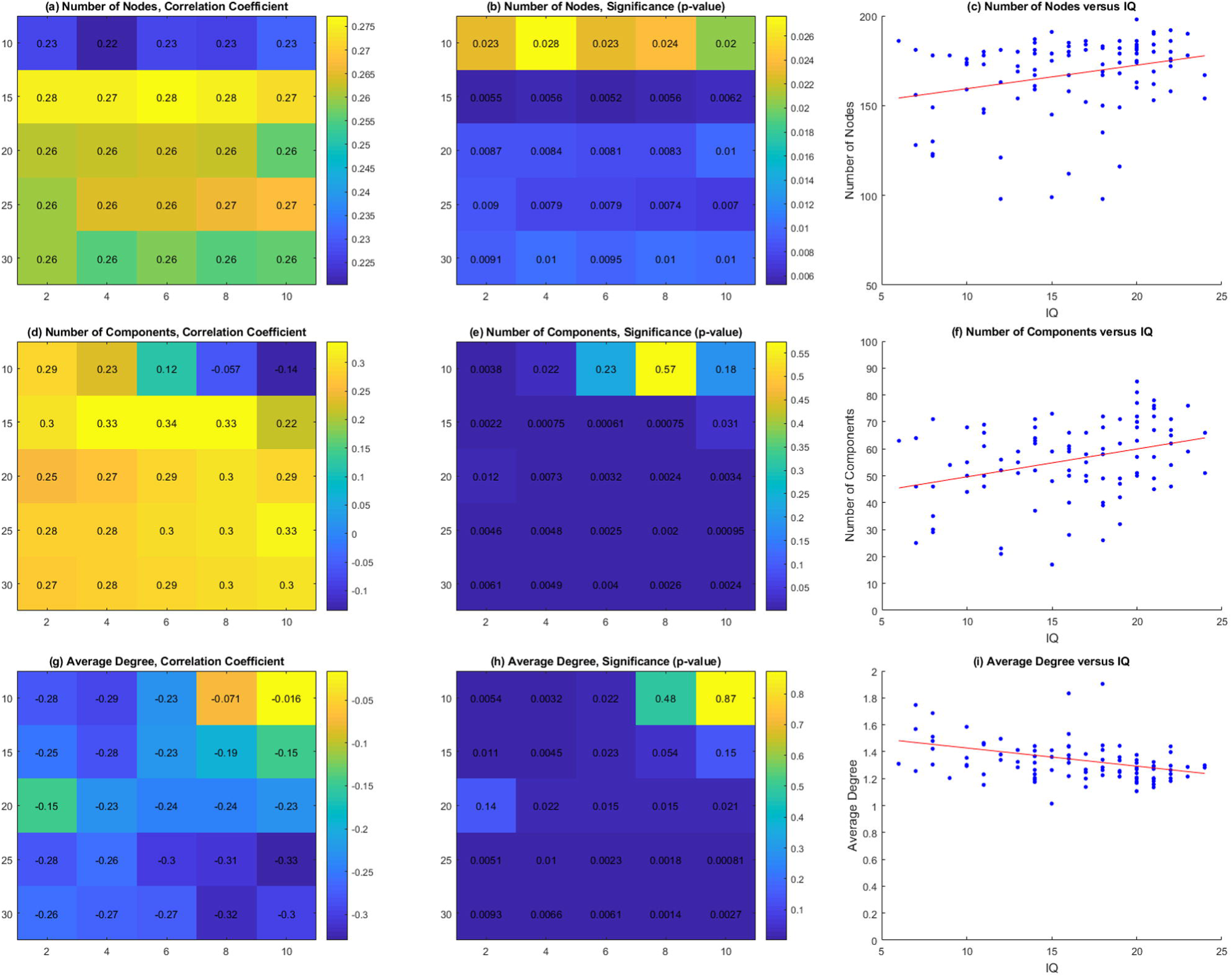
The number of nodes, number of connected components, and average degree centrality of simplicial graphs of all subjects (data is from the left-right run of the second session of HCP data-set) were calculated and then correlated with intelligence for a range of two input parameters of binning, resolution (10, 15, 20, 25, 30) and gain (2, 4, 6, 8, 10), using fixed values of two input parameters of t-SNE function, distance metric = chebychev and perplexity = 50. This was performed to show that the number of nodes and number of connected components associate positively with intelligence while the average degree centrality correlates negatively with intelligence across a wide-range of parameters. **(a) (b)** Correlation coefficients and their corresponding p-values for the number of nodes of simplicial graphs for a range of values of resolution (y axis) and gain (x axis). **(c)** The number of nodes versus intelligence (IQ) for the lowest p-value (0.0052) and highest correlation (r = 0.28). **(d) (e)** Correlation coefficients and their corresponding p-values for the number of connected components of simplicial graphs for a range of values of resolution (y axis) and gain (x axis). **(f)** The number of connected components versus intelligence (IQ) for the lowest p-value (0.00061) and highest correlation (r = 0.34). **(g) (h)** Correlation coefficients and their corresponding p-values for the average degree centrality of simplicial graphs for a range of values of resolution (y axis) and gain (x axis). **(i)** The average degree centrality versus intelligence (IQ) for the lowest p-value (0.00081) and highest absolute correlation (r = −0.33).

In addition, to further strengthen our claims, we ran the preceding systematic investigation comprised of three steps on the right-left run of the first and second sessions of HCP data-set. The corresponding results depicted in Fig. S3 (the step 1 which corresponds to Fig. 3), Fig. S4 (the step 2 which corresponds to Fig. 4), and Fig. S5 (the step 3 which corresponds to Fig. 5) generally corroborate our three main hypotheses. In these three figures, for the most range of parameters and for all significant p-values, the number of nodes associates positively with intelligence. Moreover, based on these three figures and for the most range of parameters, the number of components associates positively with intelligence; however, for the lowest p-value, this correlation is negative. Finally, based on these three figures and for the most range of parameters, the average degree centrality associates negatively with intelligence; however, for the lowest p-value, this correlation is positive.

## DISCUSSION

Relating intelligence to functional neuroimaging or finding its neural correlates in resting-state fMRI is a long-standing goal in cognitive neuroscience. We started this paper by referring to high-level cognitive models of intelligence explaining intelligent behavior and criticizing the mainstream works on this topic which are generally in the framework of network neuroscience (Bassett et al., 2018) and intend to find the neural correlates of intelligence from functional neuroimaging data without referring to the constituents of an intelligent behavior. Even in the absence of task, we could find some features of resting-state fMRI which are correlated with intelligence and can be explained by invoking the sequential mental programming of production systems, high-level cognitive models of intelligence. Based on this general framework, we hypothesized that the functional space of resting-state fMRI signals of intelligent individuals is larger and can be partitioned to more well-separated independent components with less connections. To investigate this hypothesis, the topological data analysis was utilized to reduce the dimensionality of data and to construct the simplicial graph which captures the shape of data in two-dimensions. Afterwards, three relevant graph theoretical metrics were used and correlated with intelligence. The final results - depicted in Fig. 3, Fig. 4, and Fig. 5 for a wide-range of four input parameters of the method - show that these three metrics generally follow our hypotheses: the number of nodes and number of connected components of simplicial graphs associate positively with intelligence while the average degree centrality of simplicial graphs correlates negatively with intelligence across subjects. However, there are three issues regarding our approach which are addressed in the following three paragraphs.

The topological data analysis is a powerful method to capture the shape of high-dimensional data which was introduced recently to neuroimaging literature. Its main advantage is that it is able to decipher the overall structure of fMRI data by using holistic measures instead of more traditional pairwise relationships between the signals of nodes or voxels captured by functional connectivity matrices. In task-based fMRI, the method was successfully able to find meaningful relationships between the community structure of the simplicial graph of each subject with task performance. This was captured by positive association of the graph modularity with task performance (Saggar et al., 2018). However, in resting-state fMRI - as it is depicted in Fig. 2, Fig. S1, and Fig. S2 – the simplicial graph does not have any particular shape or structure, and as a result, it is not straightforward to define relevant graph theoretical metrics which subsequently could be related to any behavioral or psychometric measures. In this paper, we could define minimally three graph theoretical metrics which were correlated with intelligence and confirmed generally our initial hypotheses.

The other relevant concern about the topological data analysis is that the method is parametric; as an instance, in our case, it accepts four input parameters (filtering parameters: distance metric and perplexity; binning parameters: resolution and gain). The depicted results in Fig. 3, Fig. 4, and Fig. 5 indicate that the correlation values and their corresponding p-values are, to some extent, sensitive to the choice of these parameters. This is also evident in the original publication (Saggar et al., 2018) in Fig. S8 and Fig. S10. Finding the best values of these parameters which describe the data objectively is a challenging task particularly in the resting-state fMRI due to not having different states. Furthermore, it is not straightforward to generalize the obtained results to other similar data-sets using the same four input parameters. In our case, we performed optimization on these four input parameters separately for the left-right and right-left runs of the first (training-set) and second (validation-set) sessions of HCP data-set. While our main hypothesis regarding the positive correlation of the number of nodes of simplicial graphs with intelligence generally hold for both runs, but still there is a need to find an objective way to optimize the values of the parameters of this method for a wide-range of similar data-sets.

Furthermore, while we believe that t-SNE as a non-linear dimensionality reduction method is a suitable starting choice for testing the applicability of topological data analysis on this kind of problem, its overall place in the topological data analysis method seems somewhat arbitrary. As a result, it makes sense to test other dimensionality reduction methods in the general pipeline of topological data analysis to further expand the validity of our claims. However, it is worth mentioning that doing this enlarges the optimization space of the free parameters and adds to the computational complexity of the problem.

## CONCLUSION

To conclude, in this paper, high-level cognitive models of intelligence - which are explaining behaviors - were used to make hypotheses regarding the relations between the features of resting-state fMRI and psychometric measures of intelligence. The main hypothesis was about the positive association of the size of functional space of resting-state fMRI with intelligence which could be interpreted as the functional equivalent of the structural relationship between brain size and intelligence asserting that people with higher intelligence have higher total brain volume (Pietschnig, Penke, Wicherts, Zeiler, & Voracek, 2015; Ritchie et al., 2015). To test this hypothesis, a recent powerful method - the topological data analysis - was applied on the resting-state fMRI for the first time. Then, three graph theoretical metrics were defined and correlated with intelligence, which subsequently corroborated our predictions. We believe our approach in considering high-level cognitive models of intelligent behavior to make hypothesis and to perform data analysis is novel and opens a door to more detailed investigation of functional space of fMRI.

## AKNOWLEDGEMENT

This work was supported in part by the Cognitive Sciences and Technologies Council of Iran under the Iran-Brazil Collaboration Desk. We would like to thank all team members of the international project on “Prediction of human intelligence through neuroimaging features,” for fruitful discussions. Also, AD would like to thank other members of the Biomedical Engineering Lab. at the University of Tehran for discussion and hospitality. He also would like to thank Alireza Rahmat-Abadi for his help with data visualization.

## SUPPLEMENTARY MATERIAL

This article has an accompanying supplementary material.

## REFERENCES

Barbey, A. K. (2018). Network Neuroscience Theory of Human Intelligence. Trends in Cognitive Sciences, 22(1), 8–20. https://doi.org/10.1016/j.tics.2017.10.001

Bassett, D. S., Zurn, P., & Gold, J. I. (2018). On the nature and use of models in network neuroscience. Nature Reviews Neuroscience. https://doi.org/10.1038/s41583-018-0038-8

Basten, U., Hilger, K., & Fiebach, C. J. (2015). Where smart brains are different: A quantitative meta-analysis of functional and structural brain imaging studies on intelligence. Intelligence, 51, 10–27. https://doi.org/10.1016/j.intell.2015.04.009

Cole, M. W., Yarkoni, T., Repovs, G., Anticevic, A., & Braver, T. S. (2012). Global Connectivity of Prefrontal Cortex Predicts Cognitive Control and Intelligence. Journal of Neuroscience, 32(26), 8988–8999. https://doi.org/10.1523/JNEUROSCI.0536-12.2012

Deary, I. J., Penke, L., & Johnson, W. (2010). The neuroscience of human intelligence differences. Nature Reviews Neuroscience, 11(3), 201–211. https://doi.org/10.1038/nrn2793

Deco, G., Jirsa, V. K., & McIntosh, A. R. (2011). Emerging concepts for the dynamical organization of resting-state activity in the brain. Nature Reviews Neuroscience, 12(1), 43–56. https://doi.org/10.1038/nrn2961

Dubois, J., Galdi, P., Paul, L. K., & Adolphs, R. (2018). A distributed brain network predicts general intelligence from resting-state human neuroimaging data. Philosophical Transactions of the Royal Society B.

Duncan, J. (2010a). How Intelligence Happens. Yale University Press.

Duncan, J. (2010b). The multiple-demand (MD) system of the primate brain: mental programs for intelligent behaviour. Trends in Cognitive Sciences, 14(4), 172–179. https://doi.org/10.1016/j.tics.2010.01.004

Duncan, J., Chylinski, D., Mitchell, D. J., & Bhandari, A. (2017). Complexity and compositionality in fluid intelligence. Proceedings of the National Academy of Sciences, 114(20), 5295–5299. https://doi.org/10.1073/pnas.1621147114

Duncan, J., Seitz, R. J., Kolodny, J., Bor, D., Herzog, H., Ahmed, A., … Emslie, H. (2000). A neural basis for general intelligence. Science, 289(5478), 457–460. https://doi.org/10.1126/science.289.5478.457

Fedorenko, E., Duncan, J., & Kanwisher, N. (2013). Broad domain generality in focal regions of frontal and parietal cortex. Proceedings of the National Academy of Sciences. https://doi.org/10.1073/pnas.1315235110

Finn, E. S., Shen, X., Scheinost, D., Rosenberg, M. D., Huang, J., Chun, M. M., … Todd Constable, R. (2015). Functional connectome fingerprinting: Identifying individuals based on patterns of brain connectivity. Nature Neuroscience, 18(11), 1664–1671. https://doi.org/10.1038/nn.4135

Genç, E., Fraenz, C., Schlüter, C., Friedrich, P., Hossiep, R., Voelkle, M. C., … Jung, R. E. (2018). Diffusion markers of dendritic density and arborization in gray matter predict differences in intelligence. Nature Communications, (2018). https://doi.org/10.1038/s41467-018-04268-8

Glasser, M. F., Sotiropoulos, S. N., Wilson, J. A., Coalson, T. S., Fischl, B., Andersson, J. L., … Hcp, W. (2013). The minimal preprocessing pipelines for the Human Connectome Project. NeuroImage, 80, 105–124. https://doi.org/10.1016/j.neuroimage.2013.04.127

Greene, A. S., Gao, S., Scheinost, D., & Constable, R. T. (2018). Task-induced brain state manipulation improves prediction of individual traits. Nature Communications, (2018). https://doi.org/10.1038/s41467-018-04920-3

Hampshire, A., Highfield, R. R., Parkin, B. L., & Owen, A. M. (2012). Fractionating Human Intelligence. Neuron, 76(6), 1225–1237. https://doi.org/10.1016/j.neuron.2012.06.022

Jung, R. E., & Haier, R. J. (2007). The Parieto-Frontal Integration Theory (P-FIT) of intelligence: Converging neuroimaging evidence. Behavioral and Brain Sciences, 30(2), 135–154. https://doi.org/10.1017/S0140525X07001185

Kriegeskorte, N., & Douglas, P. K. (2018). Cognitive computational neuroscience. Nature Neuroscience. https://doi.org/10.1038/s41593-018-0210-5

Kruschwitz, J. D., Waller, L., Daedelow, L. S., Walter, H., & Veer, I. M. (2018). General, crystallized and fluid intelligence are not associated with functional global network efficiency: A replication study with the human connectome project 1200 data set. NeuroImage, 171(January), 323–331. https://doi.org/10.1016/j.neuroimage.2018.01.018

Laland, K. N. (2018). Darwin’s Unfinished Symphony: How Culture Made the Human Mind. Princeton University Press.

Lum, P. Y., Singh, G., Lehman, A., Ishkanov, T., Alagappan, M., Carlsson, J., & Carlsson, G. (2013). Extracting insights from the shape of complex data using topology. Scientific Reports, 1–8. https://doi.org/10.1038/srep01236

Maaten, L. Van Der, & Hinton, G. (2008). Visualizing Data using t-SNE. Journal of Machine Learning Research, 9, 2579–2605.

Neubauer, A. C., & Fink, A. (2009). Intelligence and neural efficiency. Neuroscience and Biobehavioral Reviews, 33(7), 1004–1023. https://doi.org/10.1016/j.neubiorev.2009.04.001

Pietschnig, J., Penke, L., Wicherts, J. M., Zeiler, M., & Voracek, M. (2015). Meta-analysis of associations between human brain volume and intelligence differences: How strong are they and what do they mean? Neuroscience and Biobehavioral Reviews, 57, 411–432. https://doi.org/10.1016/j.neubiorev.2015.09.017

Pinker, S. (2009). How the Mind Works. W. W. Norton.

Ritchie, S. J., Booth, T., Hernández, C. V., Corley, J., Muñoz, S., Gow, A. J., … Deary, I. J. (2015). Beyond a bigger brain: Multivariable structural brain imaging and intelligence. Intelligence, 51, 47–56. https://doi.org/10.1016/j.intell.2015.05.001

Saggar, M., Sporns, O., Gonzalez-castillo, J., Bandettini, P. A., Carlsson, G., Glover, G., & Reiss, A. L. (2018). Towards a new approach to reveal dynamical organization of the brain using topological data analysis. Nature Communications, (2018). https://doi.org/10.1038/s41467-018-03664-4

Schultz, X. D. H., & Cole, X. W. (2016). Higher Intelligence Is Associated with Less Task-Related Brain Network Reconfiguration. Journal of Neuroscience, 36(33), 8551–8561. https://doi.org/10.1523/JNEUROSCI.0358-16.2016

Shen, X., Tokoglu, F., Papademetris, X., & Constable, R. T. (2013). Groupwise whole-brain parcellation from resting-state fMRI data for network node identification. NeuroImage, 82, 403–415. https://doi.org/10.1016/j.neuroimage.2013.05.081

Song, M., Zhou, Y., Li, J., Liu, Y., Tian, L., Yu, C., & Jiang, T. (2008). Brain spontaneous functional connectivity and intelligence. NeuroImage, 41(3), 1168–1176. https://doi.org/10.1016/j.neuroimage.2008.02.036

Tschentscher, N., Mitchell, X. D., & Duncan, X. J. (2017). Fluid Intelligence Predicts Novel Rule Implementation in a Distributed Frontoparietal Control Network. Journal of Neuroscience, 37(18), 4841–4847. https://doi.org/10.1523/JNEUROSCI.2478-16.2017

van den Heuvel, M. P., & Hulshoff Pol, H. E. (2010). Exploring the brain network: A review on resting-state fMRI functional connectivity. European Neuropsychopharmacology, 20(8), 519– 534. https://doi.org/10.1016/j.euroneuro.2010.03.008

van den Heuvel, M. P., Stam, C. J., Kahn, R. S., & Hulshoff Pol, H. E. (2009). Efficiency of Functional Brain Networks and Intellectual Performance. Journal of Neuroscience, 29(23), 7619–7624. https://doi.org/10.1523/JNEUROSCI.1443-09.2009

