## Supplementary Material for "Resting-state fMRI Signals of Intelligent People Wander in a Larger Space"

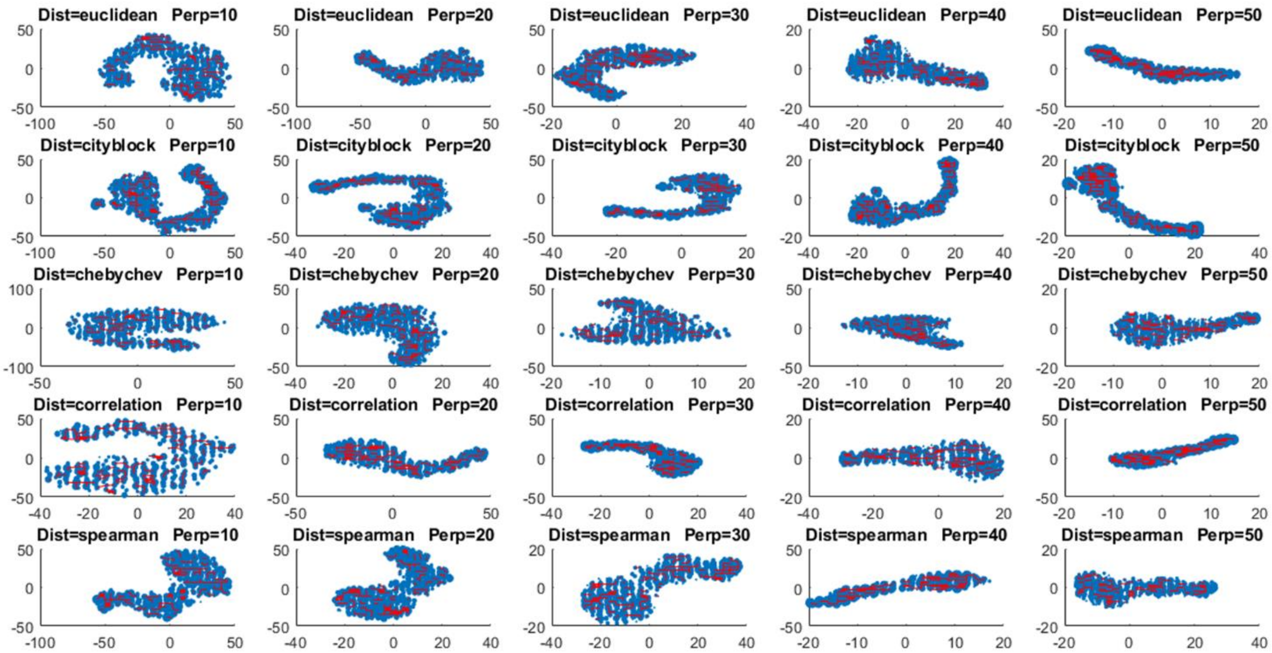


**Fig. S1:** Two input parameters of filtering function, t-SNE, were purturbed while using the fixed values for two input parameters of binning (resolution = 20 and gain = 6) to see their effects on the results of the topological data analysis (the data for these plots belongs to a representative subject = 100307 in the left-right run of the first session of HCP data-set). For measuring distances, the following metrics were used: euclidean, cityblock, chebychev, correlation, and spearman. Also, the perplexity values were set to the following quantities: 10, 20, 30 , 40, and 50. As it is evident, the shape of the simplicial graph changes considerably across values of both parameters but it does not have any particular structure.

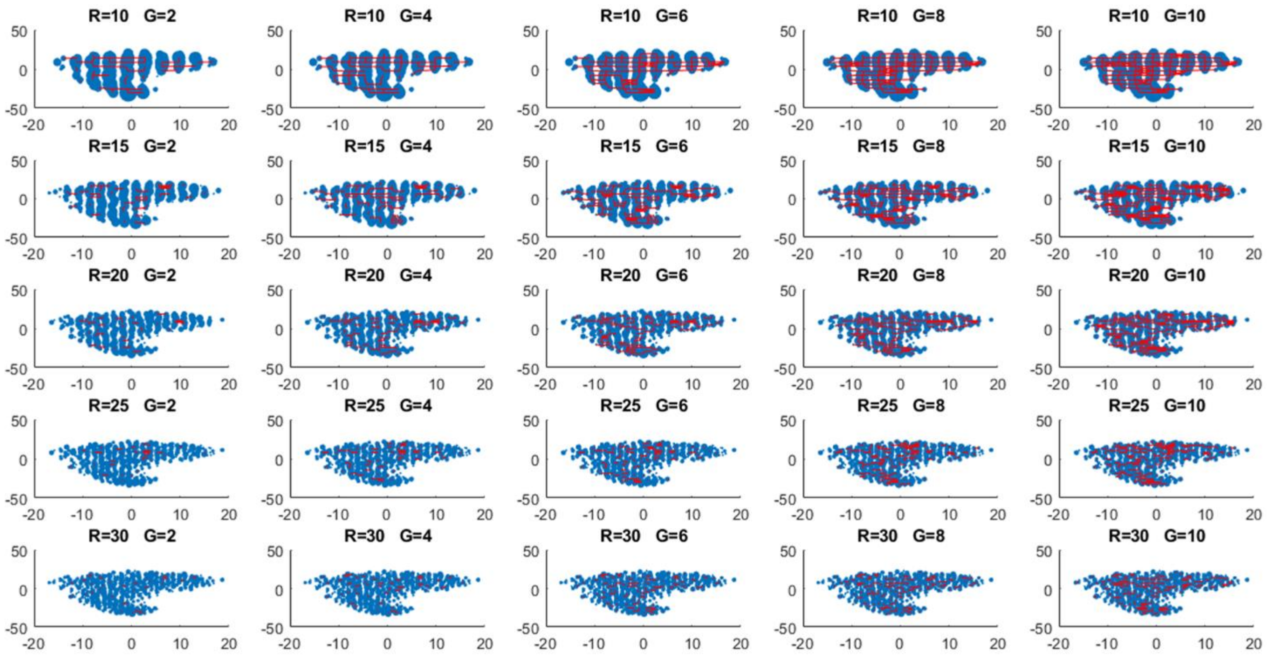


**Fig. S2:** Two input parameters of binning were purturbed while using the fixed values for two input parameters of filtering function, t-SNE (distance metric = chebychev and perplexity = 30), to see their effects on the results of the topological data analysis (the data for these plots belongs to a representative subject = 100307 in the left-right run of the first session of HCP data-set). For resolution, the following values were used: 10, 15, 20, 25, and 30. Also, the values for gain were set to the following quantities: 2, 4, 6 , 8, and 10. As it is evident, the shape of the simplicial graph changes considerably across values of both parameters but it does not have any particular structure.

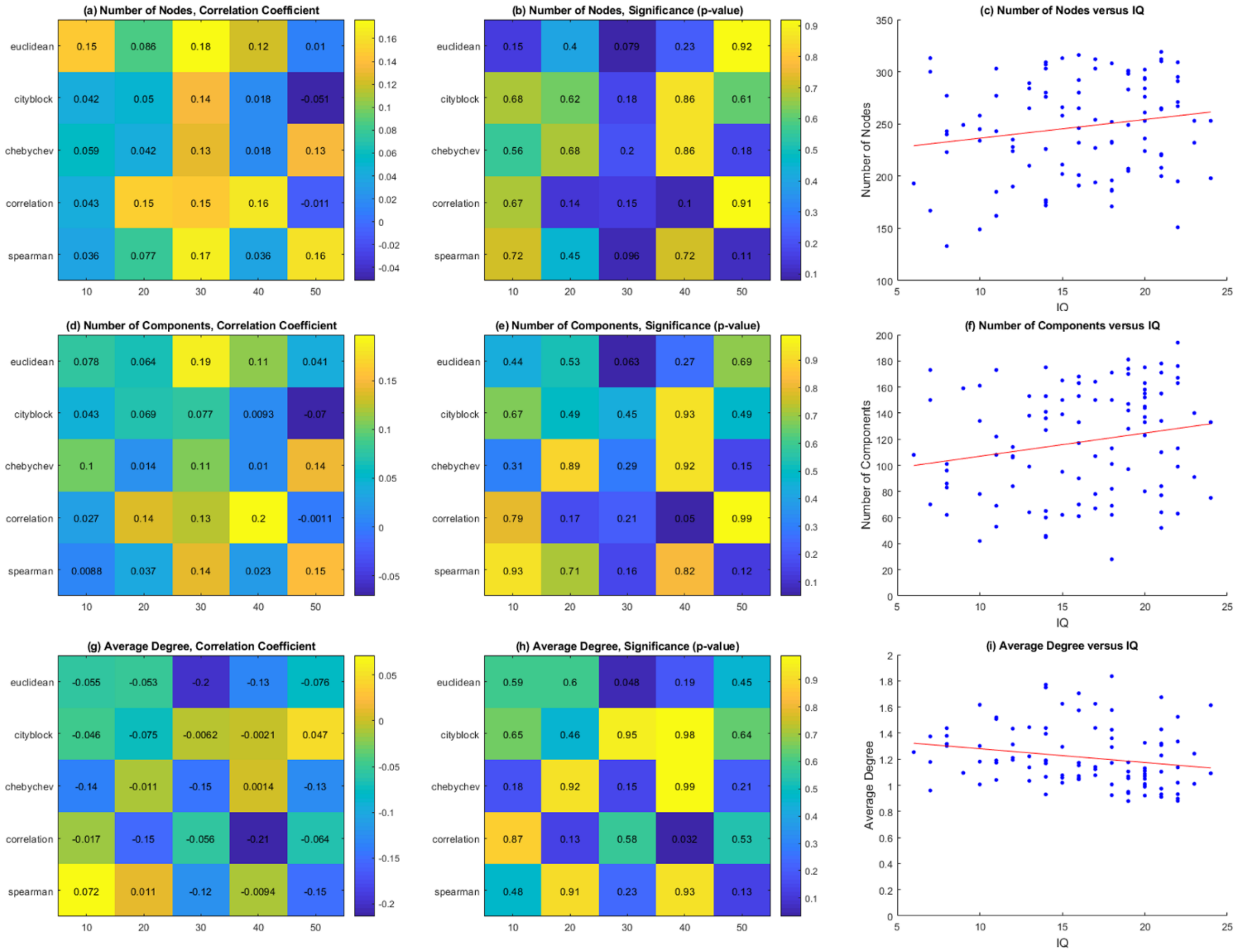


**Fig. S3:** The number of nodes, number of connected components, and average degree centrality of simplicial graphs of all subjects (data is from the right-left run of the first session of HCP data-set) were calculated and then correlated with intelligence for a range of two input parameters of t-SNE function, distance metric (euclidean, cityblock, chebychev, correlation, spearman) and perplexity (10, 20, 30, 40, 50), using fixed values of two input parameters of binning, resolution = 20 and gain = 6. This was performed to show that the number of nodes and number of connected components associate positively with intelligence while the average degree centrality correlates negatively with intelligence across a wide-range of parameters. **(a) (b)** Correlation coefficients and their corresponding p-values for the number of nodes of simplicial graphs for a range of values of distance metric (y axis) and perplexity (x axis). **(c)** The number of nodes versus intelligence (IQ) for the lowest p-value (0.079) and highest correlation (r = 0.18). **(d) (e)** Correlation coefficients and their corresponding p-values for the number of connected components of simplicial graphs for a range of values of distance metric (y axis) and perplexity (x axis). **(f)** The number of connected components versus intelligence (IQ) for the lowest p-value (0.05) and highest correlation (r = 0.2). **(g) (h)** Correlation coefficients and their corresponding p-values for the average degree centrality of simplicial graphs for a range of values of distance metric (y axis) and perplexity (x axis). **(i)** The average degree centrality versus intelligence (IQ) for the lowest p-value (0.032) and highest absolute correlation (r = - 0.21).

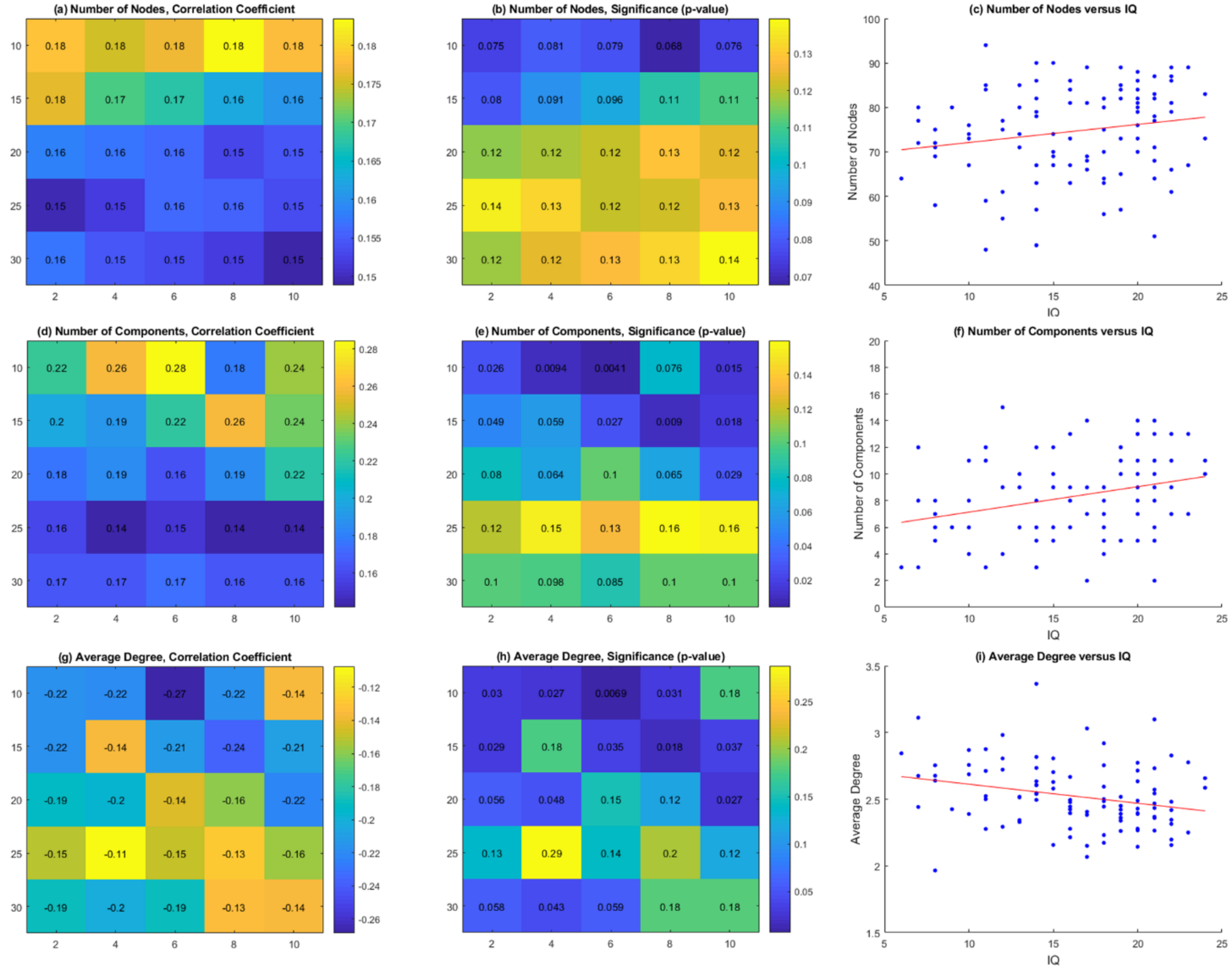


**Fig. S4:** The number of nodes, number of connected components, and average degree centrality of simplicial graphs of all subjects (data is from the right-left run of the first session of HCP data-set) were calculated and then correlated with intelligence for a range of two input parameters of binning, resolution (10, 15, 20, 25, 30) and gain (2, 4, 6, 8, 10), using fixed values of two input parameters of t-SNE function, distance metric = spearman and perplexity = 30. This was performed to show that the number of nodes and number of connected components associate positively with intelligence while the average degree centrality correlates negatively with intelligence across a wide-range of parameters. **(a) (b)** Correlation coefficients and their corresponding p-values for the number of nodes of simplicial graphs for a range of values of resolution (y axis) and gain (x axis). **(c)** The number of nodes versus intelligence (IQ) for the lowest p-value (0.068) and highest correlation (r = 0.18). **(d) (e)** Correlation coefficients and their corresponding p-values for the number of connected components of simplicial graphs for a range of values of resolution (y axis) and gain (x axis). **(f)** The number of connected components versus intelligence (IQ) for the lowest p-value (0.0041) and highest correlation (r = 0.28). **(g) (h)** Correlation coefficients and their corresponding p-values for the average degree centrality of simplicial graphs for a range of values of resolution (y axis) and gain (x axis). **(i)** The average degree centrality versus intelligence (IQ) for the lowest p-value (0.0069) and highest absolute correlation (r = - 0.27).

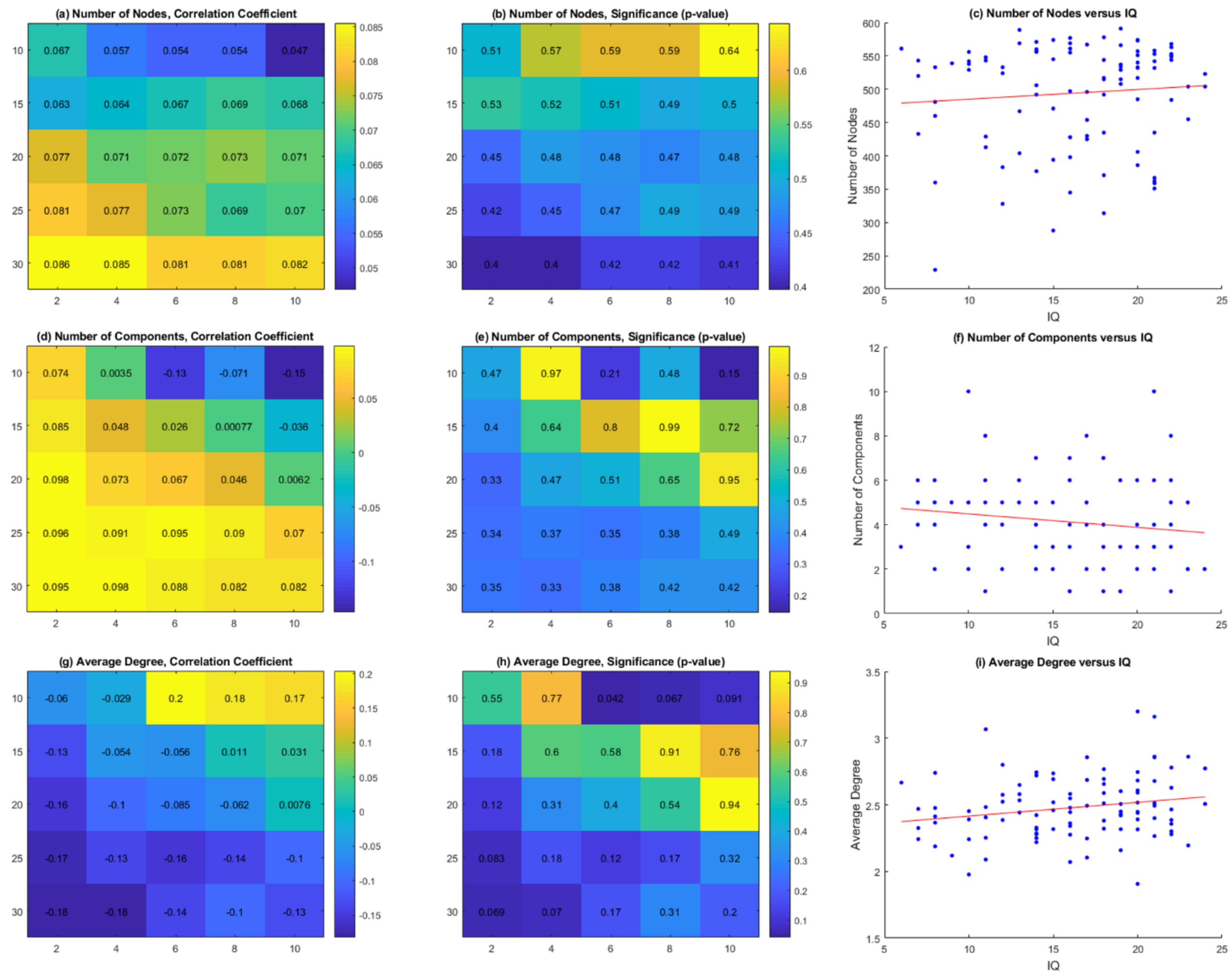


**Fig. S5:** The number of nodes, number of connected components, and average degree centrality of simplicial graphs of all subjects (data is from the right-left run of the second session of HCP data-set) were calculated and then correlated with intelligence for a range of two input parameters of binning, resolution (10, 15, 20, 25, 30) and gain (2, 4, 6, 8, 10), using fixed values of two input parameters of t-SNE function, distance metric = spearman and perplexity = 30. For all parameters, we could show that the number of nodes associates positively with intelligence albeit with not significant p-values. Also, for most of parameters, we could show that the number of connected components associates positively with intelligence; however, the lowest p-value is for a negative correlation. Finally, for most parameters, we could show that the average degree centrality associates negatively with intelligence; however, the lowest p-value is for a positive correlation. **(a) (b)** Correlation coefficients and their corresponding p-values for the number of nodes of simplicial graphs for a range of values of resolution (y axis) and gain (x axis). **(c)** The number of nodes versus intelligence (IQ) for the lowest p-value (0.4) and highest correlation (r = 0.085). **(d) (e)** Correlation coefficients and their corresponding p-values for the number of connected components of simplicial graphs for a range of values of resolution (y axis) and gain (x axis). **(f)** The number of connected components versus intelligence (IQ) for the lowest p-value (0.15) and highest absolute correlation (r = - 0.15). **(g) (h)** Correlation coefficients and their corresponding p-values for the average degree centrality of simplicial graphs for a range of values of resolution (y axis) and gain (x axis). **(i)** The average degree centrality versus intelligence (IQ) for the lowest p-value (0.042) and highest correlation (r = 0.2).
